# Adaptive therapy achieves long-term control of chemotherapy resistance in high grade ovarian cancer

**DOI:** 10.1101/2023.07.21.549688

**Authors:** Helen Hockings, Eszter Lakatos, Weini Huang, Maximilian Mossner, Mohammed Ateeb Khan, Stephen Metcalf, Francesco Nicolini, Kane Smith, Ann-Marie Baker, Trevor A. Graham, Michelle Lockley

## Abstract

Drug resistance results in poor outcomes for most patients with metastatic cancer. Adaptive Therapy (AT) proposes to address this by exploiting presumed fitness costs incurred by drug-resistant cells when drug is absent, and prescribing dose reductions to allow fitter, sensitive cells to re-grow and re-sensitise the tumour. However, empirical evidence for treatment-induced fitness change is lacking. We show that fitness costs in chemotherapy-resistant ovarian cancer cause selective decline and apoptosis of resistant populations in low-resource conditions. Moreover, carboplatin AT caused fluctuations in sensitive/resistant tumour population size *in vitro* and significantly extended survival of tumour-bearing mice. In sequential blood-derived cell-free DNA and tumour samples obtained longitudinally from ovarian cancer patients during treatment, we inferred resistant cancer cell population size through therapy and observed it correlated strongly with disease burden. These data have enabled us to launch a multicentre, phase 2 randomised controlled trial (ACTOv) to evaluate AT in ovarian cancer.

## Introduction

Systemic cancer treatment has been based on the principle that delivery of high drug doses will increase the likelihood of eradicating all malignant cells and achieve cure. Unfortunately this paradigm frequently fails, especially in the management of advanced and metastatic solid cancers, regardless of tumour type or specific drug therapy^1^. This is likely a direct consequence of therapy selecting for pre-existing drug-resistant subclones that arise stochastically due to the large number of somatic mutations that accumulate during tumour expansion^2^ or could be due to the inherent plasticity in cancer cell phenotype^3^. The paucity of available anti-cancer drug therapies means that emergence of a sufficiently large resistant cancer cell population will eventually result in treatment-resistant cancers^1^.

Evolutionary theory states that treatment-resistance should come at a fitness cost^4^ that becomes apparent when the cancer cell is in an environment that exposes the cost. Trait evolution is usually subject to tradeoffs^5^; if a cancer clone evolves to become optimal at a particular trait, such as maintaining a resistant phenotype, it will inevitably come at the price of being less good at another, for example proliferation, metastasis or invasion^6, 7^. It follows that the relative fitness of drug-sensitive and -resistant cells is reversed by drug therapy, such that in the presence of drug, resistant cells are fitter, whereas in the absence of drug sensitive cells have higher fitness^8^. Similar to other biological systems, the tradeoff between reproduction and survival is expected to be most apparent in low-resource settings where the limited resources expose “suboptimal” phenotypes. Adaptive therapy (AT) is a new treatment paradigm that exploits these competitive interactions between sensitive and resistant subclones^7^, aiming to maintain a sufficient population of sensitive cells to suppress the proliferation of ‘less fit’ resistant cells^9^. This approach accepts that within the palliative setting, cancer cannot be eradicated and aims to control rather than cure^8, 10^.

Adaptive therapy is predicted to be beneficial in mathematical models^11, 12^ and has been shown to prolong survival in preclinical *in vivo* models^4, 13^. The first AT trial to report outcomes used the oral CYP17A1 androgen synthesis inhibitor abiraterone in metastatic castration-resistant prostate cancer^14, 15^. Abiraterone AT was directed by changes in the serum tumour marker PSA (prostate-specific antigen) as a proxy for total tumour burden. Men who received abiraterone AT experienced a prolonged median time to progression of 33.5 months compared to 14.3 months in the non-randomised control group of contemporaneously treated men who received standard continuous daily dosing (*p*<0.001). Median overall survival was also increased by AT (58.5 *vs.* 31.3 months, hazard ratio (HR) 0.41, 95% confidence interval (CI) 0.20–0.83, *p*<0.001). Importantly, AT-treated men received no abiraterone for 46% of the time on trial, with associated benefits in terms of treatment-related side effects and healthcare costs. A major criticism of this trial is that it was not randomised and compared outcomes to a contemporaneous cohort. A larger, randomised trial is currently recruiting^16^. In addition, there was no attempt to track the emergence of resistance in individual patients overtime, and so the biological mechanisms that mediated increased survival with AT remain unknown.

High-grade serous ovarian cancer (HGSC) is the most common ovarian cancer subtype, accounting for 70% of cases. Standard treatment consists of cytoreductive surgery with platinum- and taxane-based chemotherapy^17^. The recent introduction of PARP inhibitor (PARPi) maintenance therapy significantly extends progression-free survival (PFS), especially in *BRCA*-mutant and homologous recombination deficient cohorts^18–22^, and is now standard of care for all patients with advanced disease following platinum response^17^. Despite this intensive treatment, the majority of HGSC cases recur and patients are subsequently treated with multiple lines of platinum-containing chemotherapy. At each sequential relapse, chemotherapy is less effective, ultimately leading to treatment failure^23^.

This relapsing-remitting nature and clear evolution of resistance to platinum-based therapy makes HGSC an exemplary disease in which to study AT. Here we show that platinum-resistant HGSC cells exhibit reduced fitness in the absence of drug that is exposed by low resource conditions. This results in resistant population decline that is mediated by apoptosis rather than secreted factors. Our experiments reveal that this reduced fitness of resistant cells is reversed by platinum treatment such that the growth dynamics of sensitive/resistant populations fluctuate during drug therapy. Importantly, we demonstrate a significant advantage of carboplatin AT compared to standard dosing in mice with established tumours. These discoveries have led to the multicentre, randomised ACTOv clinical trial (<u>A</u>daptive <u>C</u>hemo<u>T</u>herapy in <u>Ov</u>arian Cancer)^24^ that opened to recruitment in March 2023.

Like the abiraterone trial, AT dosing in ACTOv will be directed by changes in tumour burden, indicated by the serum tumour marker, cancer antigen 125 (CA125). Ideally, AT would be directed by the size of the emergent resistant population but platinum resistance is not associated with easily trackable markers such as recurrent single nucleotide variants^25, 26^ or common copy number drivers^27^. However, post-treatment HGSCs do carry new copy number alterations (CNAs) in addition to their already highly altered genomes^28^ that appear to be patient-specific. We recently developed a new bioinformatics pipeline, liquidCNA (LiqCNA)^29^, which exploits these copy number changes to quantify the emergence of treatment-resistance. Thus in situations like platinum chemotherapy, where there is no known recurrent mutational driver of resistance, LiqCNA could track the resistant population and decipher whether AT does indeed work by controlling its growth. In the current study, we use sequential blood and biopsy samples to demonstrate that LiqCNA correlates with tumour burden in HGSC patients. This crucial development is expected to improve AT by enabling the evolution of therapy resistance, rather than proxy measures of tumour burden, to direct adaptive drug dosing for future cancer patients.

## Results

### Drug-resistant HGSC cell populations exhibit a proliferative fitness cost in low resource conditions

Cisplatin- and carboplatin-resistant HGSC cells were created as we previously described^26^. In brief, OVCAR4 and Cov318 cells were cultured *in vitro* in increasing concentrations of either cisplatin or carboplatin. Dose escalation was stopped once a 2–10 fold increase in IC_50_ was achieved to reflect the resistance observed in human patients^30^. Cisplatin-resistant cells derived from OVCAR4 were denoted Ov4Cis and those derived from Cov318 were denoted CovCis. Carboplatin-resistant OVCAR4 cells were denoted Ov4Carbo. OVCAR4 and Cov318 cells were transfected with lentiviruses expressing GFP to create OVCAR4-GFP and Cov-GFP respectively. Cisplatin- and carboplatin-resistant OVCAR4 cells were transfected with lentiviruses expressing RFP to create Ov4Cis-RFP and Ov4Carbo-RFP (**Fig.1A**). Cells were immediately expanded and aliquots were frozen. Low passage cells, cultured without drug, were used in all subsequent experiments.

**Figure 1.**
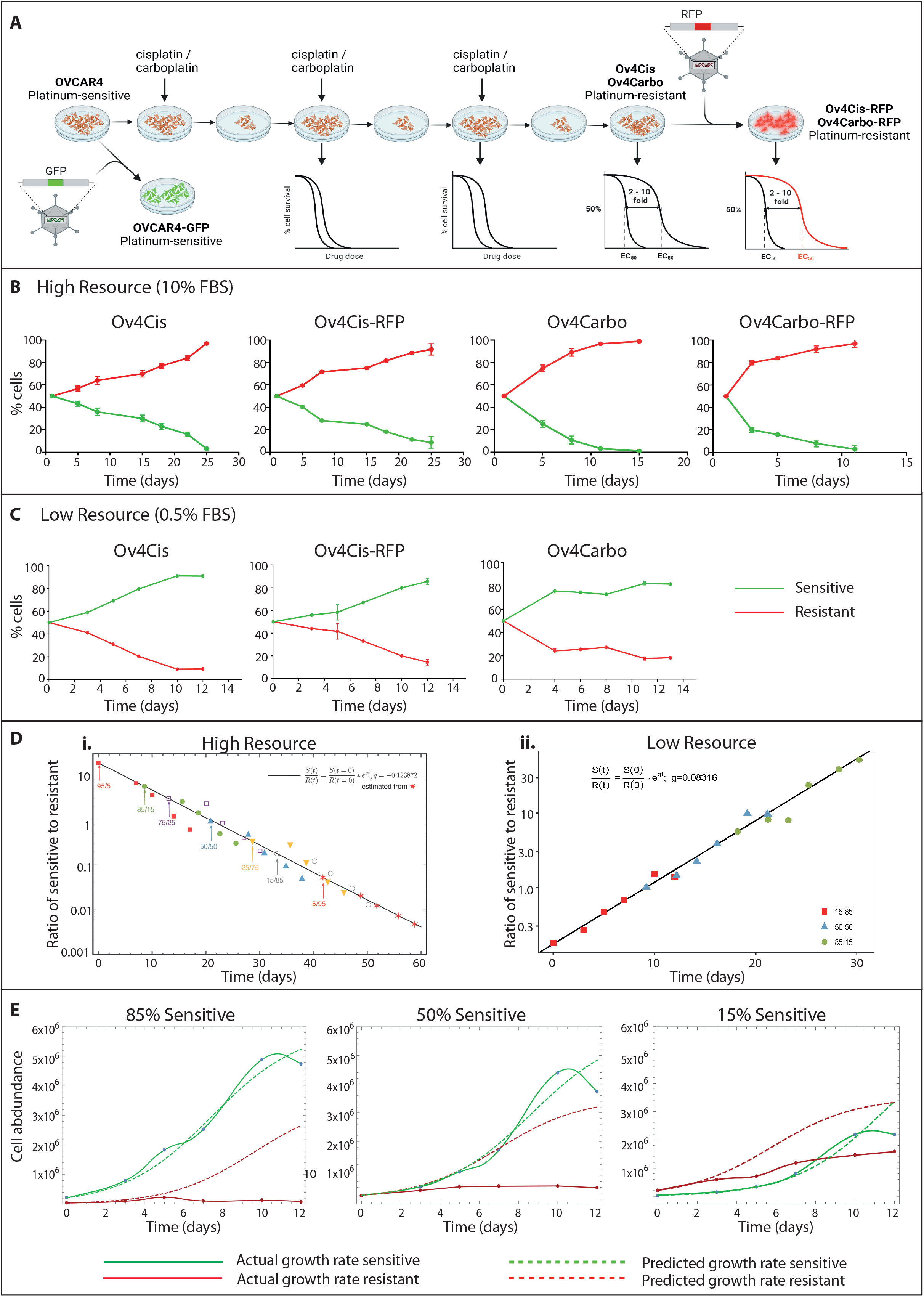
**A:** Schematic showing derivation of resistant cell panel. **B:** Abundance of sensitive (OVCAR4-GFP) and resistant (Ov4Cis, Ov4Cis-RFP, Ov4Carbo, Ov4Carbo-RFP) cells over time when co-cultured at a starting ratio of 50:50 in high resource conditions (10% FBS). Green=OVCAR4-GFP, red=resistant cell lines, mean±st.d, *n*=3 technical replicates. **C:** Abundance of resistant cell lines (Ov4Cis, Ov4Cis-RFP, Ov4Carbo) co-cultured with sensitive OVCAR4-GFP cells in low resource conditions (media exchanged with fresh 0.5% FBS-containing media every 24 hours). Green=OVCAR4-GFP, red=resistant cell lines, mean±st.d, *n*=3 technical replicates. **D:** The ratio of OVCAR4-GFP:OV4Cis plotted on a log scale over time in high (i) and low (ii) resource condition. The black line shows a linear fit of this log ratio based on the 5:95 (i) and 15:85 (ii) dataset. The slope of the line indicated on the plot corresponds to the difference in growth rate between sensitive and resistant cells: g_s_ - g_r_=g. Initial data points of other starting ratios were staggered to fit on the fitted line, but these data were not used for the linear fit (see Methods). **E:** Abundance of sensitive (green) and resistant cells (red) over time when co-cultured in low resource conditions (0.5% FBS exchanged daily) at three starting ratios (85% sensitive, 50% sensitive and 15% sensitive). Solid lines indicate observed cell growth, dashed lines indicate predicted cell growth based on 100% monoculture experiments.

*In vitro* co-cultures were created by seeding chemotherapy-sensitive and chemotherapy-resistant cells at different starting ratios. At each time point, total cell number was counted and the proportion of GFP- and RFP-expressing cells was quantified by FACS to calculate the abundance of sensitive and resistant cell populations over time. When cells were co-cultured in standard conditions including 10% FBS (high resource), the resistant cell population increased as a percentage of the total population over time. This was true in all 4 cell pairs (OVCAR4-GFP:Ov4Cis, OVCAR4-GFP:Ov4Cis-RFP, OVCAR4- GFP:Ov4Carbo and OVCAR4-GFP:Ov4Carbo-RFP) (**Fig.1B**) and at all starting sensitive:resistant ratios (**Fig.S1**). All cells grew exponentially with similar growth rates for all seeding ratios (0.31 ± 0.05 for sensitive cells and 0.43 ± 0.04 for resistant cells, mean ± SD) and by day 7, no cell line had reached logistic growth (**Fig.S2**). By using the difference between these two growth rates, ‘g’, estimated from the co-culture data of one seeding ratio (5:95), we fitted all measures starting from other ratios with an accuracy across four orders of magnitude (**Fig.1Di**). This gives strong evidence of independent growth without competition in high resource conditions.

Since resources are constrained in human cancer, co-culture experiments were repeated in constant low resource conditions by exchanging medium for fresh 0.5% FBS-containing medium every 24 hours. This time, although the overall population increased, the size of the resistant cell population decreased over time relative to the total population (**Fig.1C**). In these low resource conditions, growth rates of mixed sensitive and resistant populations were logistic indicating that populations were competing for resources and that carrying capacity constrained total population size. Resistant cell growth rates were lower than for sensitive cells (*g*_*s*_ − *g*_*r*_ = 0.08 doublings/day) and remained independent of the initial ratio of sensitive:resistant cells (**Fig.1Dii**). This demonstrates the fitness cost borne by drug resistant cells in resource poor-environments, and shows that competition for limited resources particularly penalises resistant cancer cells.

To further test this, OVCAR4-GFP and Ov4Cis populations were grown as co-cultures in low serum conditions at three different starting ratios. **Fig.1E** shows the measured abundance of GFP-positive (green) and GFP-negative (red) populations over time (solid lines) as well as the predicted growth of each cell population based on their initial seeding density and their measured growth as mono- cultures (dashed lines). In all ratios tested, the presence of resistant cells did not change the growth of the sensitive cells, but the presence of sensitive cells slowed the growth and lowered the cell abundance of resistant cells compared to their carrying capacity in mono-culture. Competition was also observed in the OVCAR4-GFP:Ov4Cis-RFP cell pair especially when starting from a large sensitive to resistant ratio (**Fig.S3A**). In the OVCAR4-GFP:Ov4Carbo pair, stronger competition was observed at carrying capacity (**Fig.S3B**). In the Cov-GFP:CovCis pair, growth rates and carrying capacities for the two cell lines were comparable in mono- and co-culture and competition always had a larger impact on the less abundant cell line (**Fig.S3C**).

### Competition in low resource conditions induces apoptosis in resistant HGSC cells

We next examined mechanisms of decreased resistant cell fitness in low-resource conditions. Cells were grown in low resource conditions (constant 0.5% FBS as before) either as 100% mono-cultures or in co-culture at a starting ratio of 85% sensitive OVCAR4-GFP:15% resistant Ov4Cis and cell cycle profiles were obtained by FACS for up to 13 days (**Fig.2A**). There was no significant change in the proportion of OVCAR4-GFP cells in any phase of the cell cycle in co-culture compared to mono-culture. In contrast, compared to mono-culture, more Ov4Cis cells in co-culture appeared in sub-G0 (Day 6, *p*=0.0095) and fewer Ov4Cis were in G2/M phase (Day 6, *p*=0.0480; Day 13, *p*=0.0296). Moreover, compared to OVCAR4-GFP cells, more Ov4Cis cells in the co-culture were in sub-G0 (Day 6, *p*=0.0111; Day 13, *p*=0.0439) and fewer Ov4Cis cells were in G2/M (Day 4, *p*=0.0466; Day 6, *p*=0.0416; Day 11, *p*=0.0415; Day 13 (*p*=0.0182).

**Figure 2.**
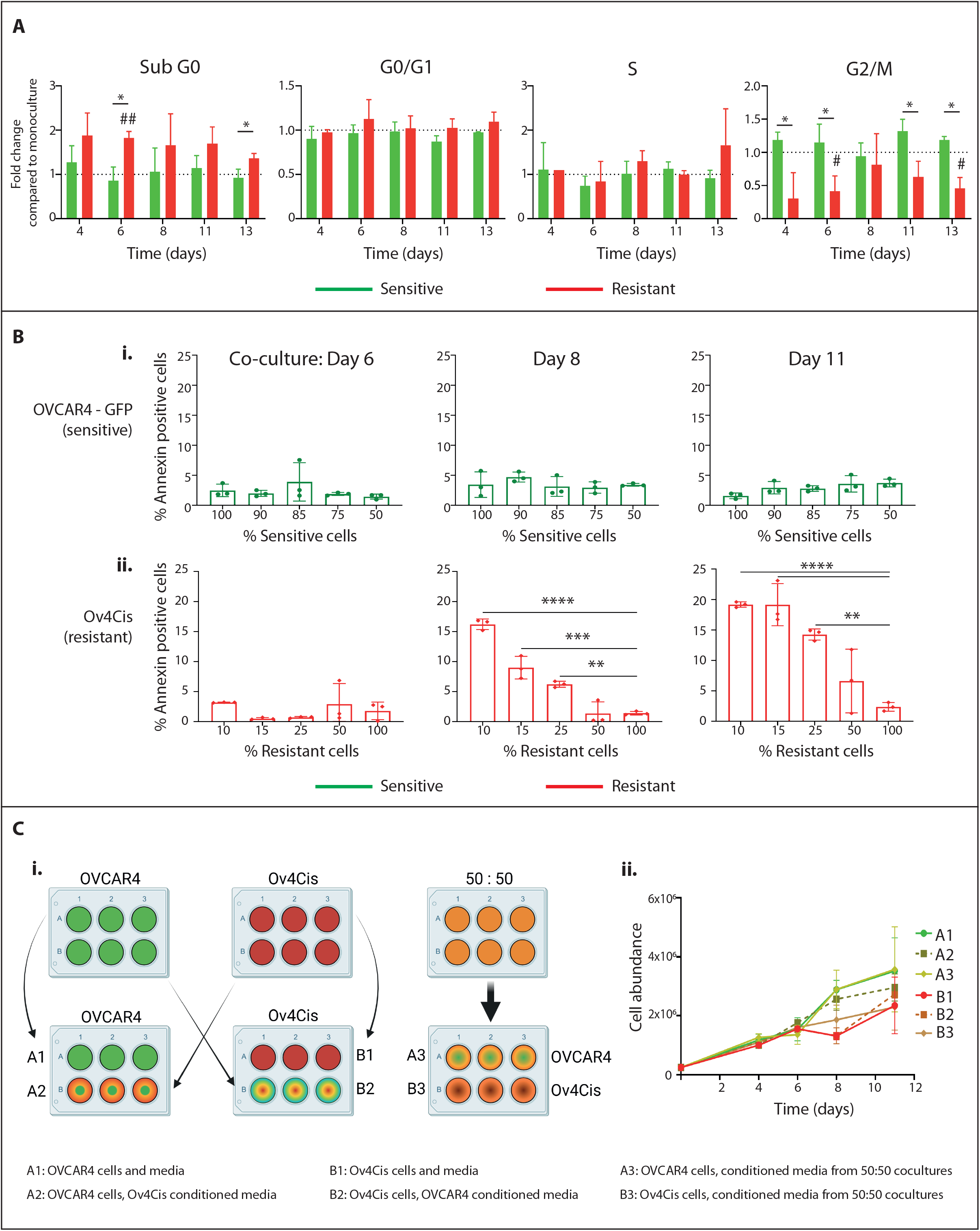
A: Cell cycle profiles of OVCAR4-GFP and Ov4Cis cells grown as mono-cultures and in co-culture (starting ratio of 85:15 OVCAR4-GFP:Ov4Cis) in constant low resource conditions (0.5% FBS exchanged daily). GFP- and RFP-positive cells are normalised to the same cell type in mono- culture at the same time point (mean±st.d, *n*=3 biological replicates). Paired *t*-test: *p<0.05 OVCAR4-GFP compared to Ov4Cis; #p<0.05, ##p<0.01 =significant difference between the same cell line in co-culture and mono-culture. **B:** OVCAR4-GFP (**i**) and Ov4Cis cells (**ii**) were grown in low resource co-culture at different starting ratios: 100:0, 90:10, 85:15, 75:25, 50:50, 0:100 and stained with PI and Annexin-V. Co-cultures were sorted into green-fluorescent (sensitive) and non- fluorescent (resistant) populations and Annexin-V positivity was measured over time by flow cytometry. Mean±st.d, *n*=3 biological replicates, 1-way ANOVA compared to 100% control samples, **p<0.01, ***p<0.001, ****p<0.0001. **Ci:** OVCAR4 (sensitive, green), Ov4Cis (resistant, red) and OVACR4:Ov4Cis 50:50 co-cultures (orange) were grown in low resource conditions. Every 24 hours, media was exchanged for media that had been pre-conditioned for 24 hours by either the same cell line (A1, B1), the other cell line (A2, B2) or a 50:50 OVCAR4:Ov4Cis co-culture (A3, B3). **Cii:** Cell abundance over time by flow cytometry (mean±st.d, *n*=3 technical replicates).

Since the sub-G0 fraction represents apoptotic cells, we repeated these low resource co-culture experiments at a range of starting ratios (OVCAR4-GFP:Ov4Cis, 100:0, 90:10, 85:15, 75:25: 50:50, 0:100) but this time added the standard apoptotic marker, Annexin-V. In sensitive OVCAR4-GFP cells, Annexin-V staining did not change over time or according to the co-culture ratio (**Fig.2Bi**). In contrast, in resistant Ov4Cis cells, Annexin-V progressively increased as the size of the sensitive population increased (**Fig.2Bii**). This was most marked at later time points (Day 8, *p*=0.0001; Day 11, *p*=0.0001 in 10% resistant co-cultures compared to 100% Ov4Cis mono-cultures) indicating that apoptosis of resistant cells contributes to their reduced abundance over time in co-culture. Sensitive (OVCAR4) and resistant (Ov4Cis) cells were then grown either as 100% mono-cultures or as 50:50 co-cultures in low resource conditions. Medium was replaced daily with conditioned medium obtained either from the same cell line, the other cell line or the 50:50 co-culture (**Fig.2Ci**) for up to 11 days. We observed no difference in growth rates or carrying capacities between the different conditions (**Fig.2Cii**), implying that the reduced growth of resistant cell populations in co-culture is not induced by secreted factors.

To investigate senescence as a possible cause of reduced growth of resistant cell populations, we again seeded co-cultures in low resource conditions, sorted cells by FACS into GFP and non-GFP populations and performed quantitative reverse transcription polymerase chain reaction (qRT-PCR) for the standard senescence markers p16 and p21. Although doxorubicin induced p16 (*p*=0.9338) and p21 (*p*=0.0201) expression in Ov4Cis cells (**Fig.S4A**), low resource co-cultures failed to induce p16 (**Fig.S4B**) or p21 (**Fig.S4C**) in either cell line for up to 13 days.

### Fitness costs of resistance *in vivo* mirror those seen in low resource conditions *in vitro*

We then used a mouse model of subcutaneous tumour growth to test if *in vivo* conditions mirrored the low resource *in vitro* setting. We mixed sensitive OVCAR4-GFP and resistant Ov4Cis-RFP cells *ex vivo* at different ratios (sensitive:resistant 100:0, 80:20, 50:50, 10:90 and 100:0) and created subcutaneous (sc) tumours by injecting mice with the same cell mixture in both flanks (*n*=2-4 mice per starting ratio). Mice were culled up to 12 weeks post injection and tumours harvested (**Fig.3A**). qPCR analysis of GFP and RFP for the sensitive and resistance cell populations respectively showed that, in all tumours, the size of the sensitive population exceeded the initial injected ratio, while the resistant population was lower than the starting ratio (**Fig.3Bi**), indicating preferential growth of the sensitive population *in vivo*. We repeated the experiment using a more clinically relevant intraperitoneal (IP) model by injecting mixtures of sensitive (OVCAR4, non-fluorescent) and resistant (Ov4Cis-RFP) cells IP (sensitive:resistant, 100:0, 80:20, 50:50, 10:90 and 100:0, *n*=3-5 mice per group). Again, we found that the proportion of RFP-positive resistant cells had again declined when measured at necropsy, 12 weeks after initial IP injection (**Fig.3Bii**).

**Figure 3.**
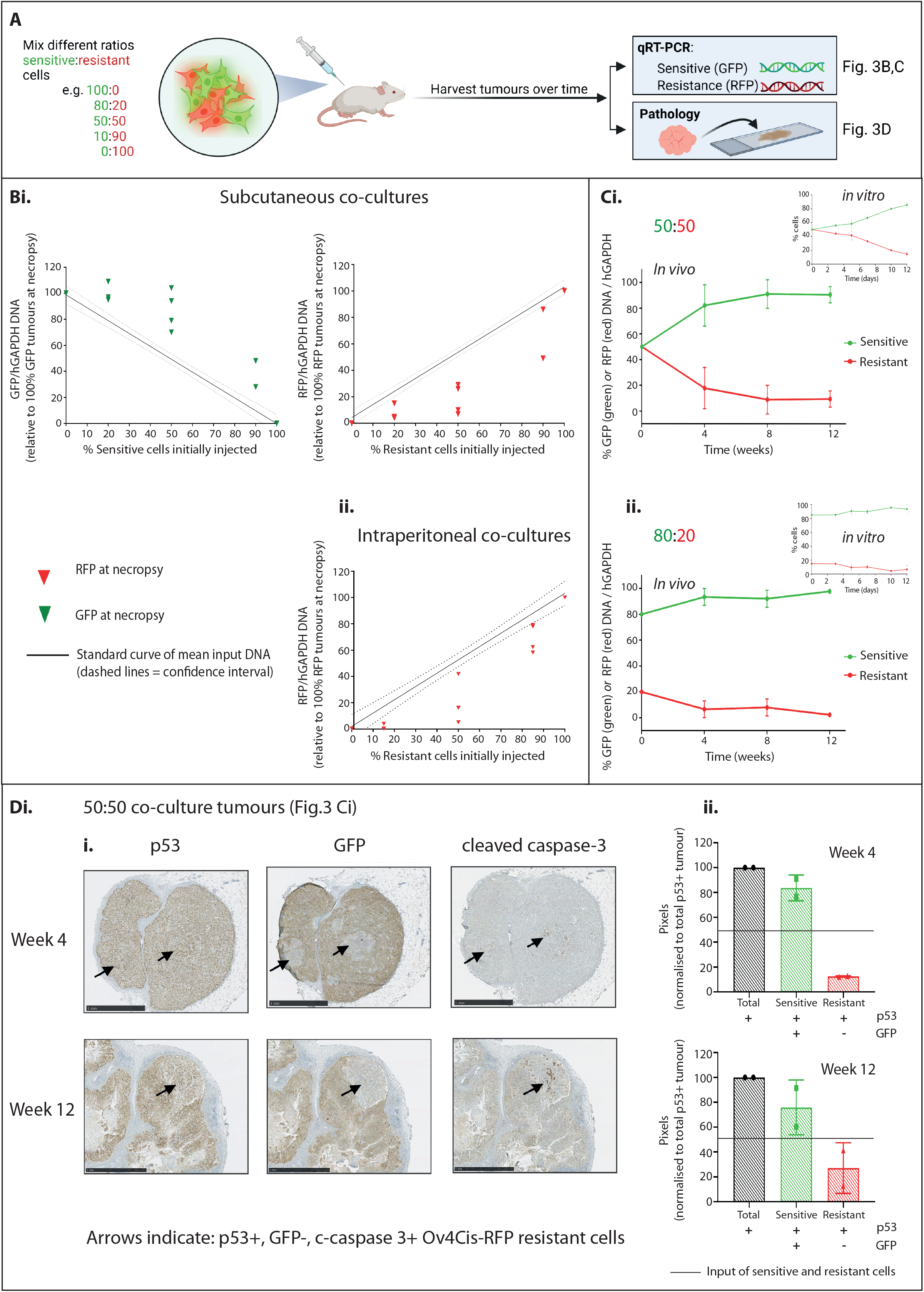
A: Schematic depicting *in vivo* co-culture experiments. **Bi:** qPCR of GFP and RFP DNA in each subcutaneous tumour co-culture (OVCAR4-GFP:Ov4Cis-RFP) and **ii:** qPCR of RFP DNA in intraperitoneal co-cultures (OVCAR4:Ov4Cis-RFP). In all cases, tumours were harvested after 12 weeks, plotted against a standard curve of input DNA (solid line=mean, dashed line=standard deviation, one icon per tumour, *n*=2-4 tumours per starting ratio). **C:** Mice were inoculated with OVCAR4-GFP and Ov4Cis-RFP cells at a ratio of either 50:50 (**i**) or 80:20 (**ii**) and culled over time. GFP and RFP DNA was measured by qPCR in each tumour, normalised to total tumour DNA (hGAPDH) and relative to 100% GFP and 100% RFP tumours (mean±st.d, *n*=4 tumours per condition). Figure insets show the same cell pair and stating ratio as *in vitro* co-cultures (expanded in Supplementary Figure S6). **D:** Immunohistochemistry for p53, GFP and cleaved caspase-3 in 50:50 OVCAR4-GFP:Ov4Cis-RFP tumours harvested at weeks 4 and 12 (Fig.3Ci). **Di:** Images of representative slides; arrows indicate p53+, GFP-ve, cleaved-caspase-3+ resistant cells. **Dii:** Pixel quantification of sensitive and resistant cells, mean±st.d, *n*=2 per time point. Solid line indicates starting ratio of sensitive:resistant cells.

To characterise the temporal dynamics of resistant and sensitive cell populations *in vivo*, mice were injected subcutaneously in both flanks with mixtures of OVCAR4-GFP and Ov4Cis-RFP at a starting ratio of either 50:50 (**Fig.3Ci**) or 80:20 (**Fig.3Cii**). Four mice in each group were culled at weeks 4, 8 and 12. The ratio of sensitive and resistant populations was quantified by qPCR as before. At all time points and both starting ratios, GFP DNA increased and RFP DNA decreased compared to week 0. This preferential growth of the sensitive population was already observed at the first time point (week 4) and stabilised thereafter. These dynamics mirrored the relative growth observed when the same two cell lines were grown as *in vitro* co-cultures in limited resources (**Fig.3C** insets and **Fig.S5**).

The contralateral flank tumours from the mice with 50:50 OVCAR4-GFP:Ov4Cis-RFP tumours (shown in **Fig.3Ci**) were stained by IHC for p53 to indicate total tumour burden and GFP to indicate platinum-sensitive cells. Small, discrete islands of GFP-negative resistant cells were observed embedded within predominantly GFP-expressing tumour nodules (**Fig.3Di**). Quantification of the area of these two populations again revealed that the GFP-positive sensitive population exceeded the 50% initially injected, while the GFP-negative resistant population declined (**Fig.3Dii**). Moreover, in keeping with our *in vitro* data (**Fig.2A,B**), positivity for the apoptotic marker cleaved caspase-3 was only observed in the resistant, GFP-negative cells and this became more apparent after 12 weeks compared to 4 weeks of *in vivo* co-culture (**Fig.3Di**). The spatial contiguity of GFP-negative resistant cells implied that they were a clonal expansion, not a conglomerate of surviving cells. Together, the reduced fitness of resistant cells observed in these mouse experiments indicate that *in vivo* tumour conditions are likely resource poor and expose resistance-associated fitness costs.

### Adaptive therapy with carboplatin significantly extends survival in tumour-bearing mice

We then sought to determine whether the relative fitness of sensitive and resistant cells is altered by chemotherapy. First we investigated the effect of platinum treatment in 50:50 co-cultures of OVCAR4- GFP:Ov4Cis cells *in vitro* (**Fig.4A**). Media was changed to 0.5% FBS-containing media 24 hours after initial plating and every 24 hours thereafter as before. Cells were treated with a single dose of carboplatin (0-1μM) on D6, which was washed off after 24 hours during the next scheduled exchange of low serum media. Plates were harvested over time and the relative proportion of GFP- and RFP-positive cells was measured by FACS. As before, sensitive cell growth exceeded that of resistant cells in low resource co-cultures. Drug effect occurred several days after exposure (range: 5-12 days) and always resulted in a reduction in the proportion of sensitive cells. This relationship was dose-dependent, such that a greater reduction in the sensitive population was seen at earlier time points with higher doses of cisplatin . In all cases, sensitive cells subsequently outgrew resistant ones, presumably as the drug effect wore off, demonstrating that the size of sensitive and resistant populations changes dynamically during treatment.

**Figure 4.**
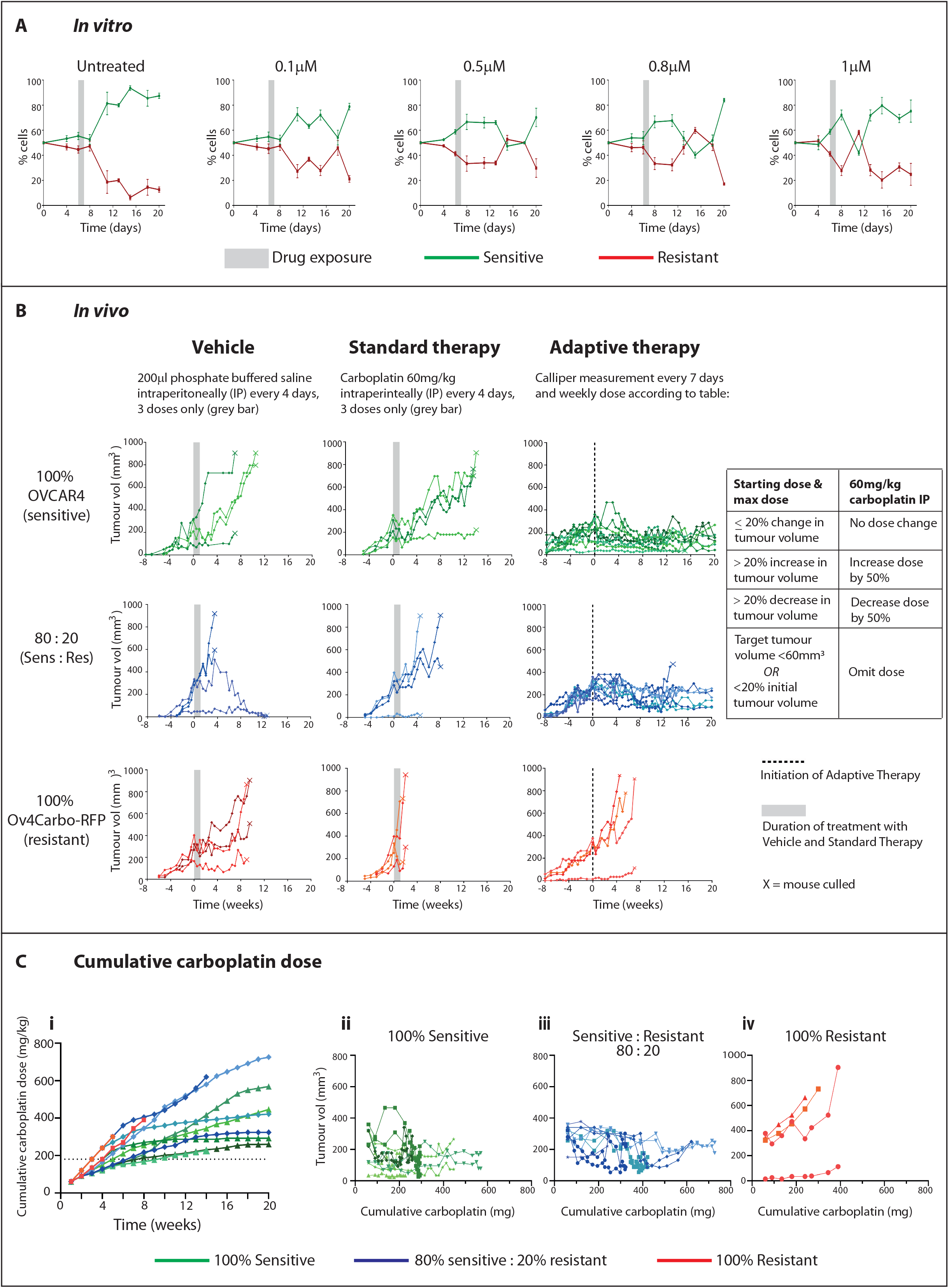
A: OVCAR4-GFP and Ov4Cis cells were grown as 100% monocultures or 50:50 co-cultures. On day 6, cells were treated with cisplatin (0.1–1µM) or vehicle (dashed line). Cultures were harvested over time and the ratio of GFP positive (green) and negative (red) cells measured by flow cytometry over time. (mean±st.d, *n*=3 technical replicates). **B:** Mice were injected subcutaneously in both flanks with 100% OVCAR4 cells (sensitive-green, top row), Ov4Carbo-RFP cells (resistant-red, bottom row) or 80:20 OVCAR4:Ov4Carbo-RFP (blue, middle row). Mice were randomised to receive (vehicle IP every four days x3; left column, *n*=2 mice per ratio), Standard Therapy (60mg/kg carboplatin IP every 4 days x3; middle column, *n*=2 mice per ratio) or Adaptive Therapy as per Table (right column *n*=3-5 mice per ratio). Grey bars indicate duration of vehicle and Standard Therapy, dotted line = initiation of Adaptive Therapy at Time 0. Each line indicates one tumour, X indicates mouse culled. **Ci:** Cumulative dose of carboplatin in mg/kg over time for all mice receiving adaptive therapy. Green=100% sensitive, blue=80% sensitive, red=100% resistant with a different shade for each mouse. **ii-iv:** Cumulative carboplatin (mg) plotted against volume of individual tumours separated into **ii:** 100% sensitive-green, **iii:** 80% sensitive– blue and **iv:** 100% resistant–red. Each line indicates one tumour and the same colour shade is used for the mouse in **Ci.**

We then sought to test Adaptive Therapy (AT) *in vivo* (**Fig.4B**) by creating subcutaneous tumours in opposing flanks of mice with either 100% OVCAR4 (sensitive) cells, 100% Ov4Carbo-RFP (carboplatin-resistant) or an 80:20 co-culture OVCAR4:Ov4Carbo-RFP. Tumours were measured weekly with callipers and for each animal, when one flank tumour reached 300mm^3^ in volume (target tumour, Time 0), that mouse was randomly allocated to one of three treatment groups: vehicle control (200µl PBS, IP every 4 days for 3 doses, *n*=2 mice per initial ratio of sensitive:resistant cells), standard carboplatin treatment (60mg/kg carboplatin IP every 4 days for 3 doses, *n*=2 mice per ratio) or carboplatin AT (IP weekly based on changes in tumour size, **Fig.4B, table,** *n*=3-5 mice per ratio).

All mice treated with vehicle and standard carboplatin reached humane endpoint before the end of the experiment (marked X, **Fig.4B**). Standard carboplatin therapy temporarily halted tumour growth in mice with majority sensitive tumours (100% OVCAR4 and 80:20 OVCAR4:Ov4Carbo-RFP) but all tumours subsequently re-grew once the treatment was stopped and there was no difference in survival between vehicle and standard carboplatin in these two tumour groups. Conversely, in mice with 100% Ov4Carbo-RFP tumours (resistant), standard carboplatin treatment reduced median survival to 5.75 weeks compared to 15.25 weeks with vehicle (*p*=0.0896).

AT significantly improved outcomes in all tumour groups. In mice with 100% Ov4Carbo-RFP tumours (resistant), AT increased survival compared to standard carboplatin (*p*=0.0389) but all mice still progressed during AT and reached humane endpoint by week 8. In mice with 100% OVCAR4 tumours (sensitive), median survival following AT was undefined because there were too few deaths in this group, compared to 18.75 weeks with standard carboplatin therapy (*p*=0.0082). One mouse in this 100% OVCAR4, AT-treated group was culled 15 weeks after treatment initiation due to unexplained weight loss in the absence of tumour progression but in the other four mice, tumour burden was controlled until experimental end point (20 weeks). In mice with 80:20 OVCAR4:Ov4Carbo-RFP tumours, AT also achieved durable tumour control such that median survival could not be defined, compared to a median survival of 11.25 weeks following standard therapy (*p*=0.574). Two of five AT treated mice with 80:20 tumours reached home office limits before experimental endpoint (four and 13.5 weeks after treatment initiation) due to large, haemorrhagic tumours, which appeared to have escaped therapeutic control (**Fig.S6A**). Histological examination revealed that the mouse that died at 4 weeks had a tumour that was dominated by resistant, luciferase-expressing cells, whereas in the mouse that was killed at 13 weeks, the enlarged tumour was cystic with a central necrotic core (**Fig.S6B**).

All AT-treated mice received a higher cumulative carboplatin dose than those in the standard therapy arm (**Fig.4Ci**). 100% resistant tumours were still able to grow despite this increase in cumulative carboplatin (**Fig.4Civ**). However, in mice with 100% (**Fig.4Cii**) or 80% (**Fig.4Ciii**) sensitive tumours, total carboplatin dose plateaued over time as tumour size was controlled, such that at later time points weekly carboplatin was either omitted completely (three mice) or repeatedly administered at very low dose (<3mg/kg, six mice) in accordance with our AT treatment protocol (**Fig.4B, table**). In AT-treated mice, no animals displayed evidence of treatment-related toxicity. AT therefore significantly improved survival compared to standard treatment and was also well-tolerated despite administration of higher cumulative carboplatin doses.

### Lineage tracing of resistant cell population using cfDNA in patients

We previously developed a bioinformatics pipeline, called LiqCNA, for estimating the size of putatively drug-resistant tumour cell populations^29^, termed the emergent resistant (ER) population. LiqCNA uses the pattern of copy number alterations (CNAs) in an evolving tumour to identify CNAs present in an emerging subclone and infer the frequency of that subclone. We used our LiqCNA algorithm to infer the evolutionary dynamics of treatment resistance in sequential blood and tissue samples from five high grade serous ovarian cancer patients during standard treatment. Three of these patients were sampled at ≥3 timepoints (patients UP0018, UP0053 and UP0055), and two patients were sampled at two timepoints (UP0042 and UP0056). We note that from only two samples, LiqCNA cannot reliably distinguish pervasive ongoing copy number (CN) instability (e.g. a new loss in a region) and measurement biases from CNAs present exclusively in a subclone. Therefore, the subclonal ratio may have been overestimated in UP0042 and UP0056.

CNA profiles showed that resistance-specific changes emerged through therapy (UP0055: **Fig.5Ai;** UP0018, UO0042, UP0053 and UP0056: **Fig.S7 Ai-Di**) and enabled quantification of an ER population in all five cases (**Fig.5Aii** and **Fig.S7 Aii-Dii**). The most prominent CNAs in the ER population are shown in **Fig.5Aiii** and **Fig.S7 Aiii-Diii** together with known oncogenic drivers and genes associated with ovarian cancer that are contained within the ER-specific CNAs. The genomic regions at which copy number changes were observed and called by the LiqCNA algorithm differed between patients, and so while they are associated with the ER population, these CNAs might not directly confer resistance.

We then compared the change in ER calculated by LiqCNA with the change in CA125 for each patient over time (**Fig.5B**). Clinical details are provided in (**Fig.5C**). In two patients (UP0018 and UP0055), LiqCNA was conducted after a short time interval during which neither patient received anti-cancer treatment. In UP0018, both samples were obtained on the same day but a larger ER population of 59.9% (53.6% - 74.0%, 95% CI) was identified in the cfDNA sample compared to 33.6% (17.3% - 54.5%, 95% CI) in the tissue sample. In patient UP0055, the tissue sample was obtained 4 weeks before the cfDNA sample but the ER estimates were comparable: cfDNA: 34.8%, (25.6% - 53.8%, 95% CI) *vs.* tissue: 42.4% (25.7% – 53.8%, 95% CI). We found a strong correlation (*R*=0.94, *p*=0.00015) between the inferred growth rate of the ER population in two sequential samples and the absolute CA125 at the time of the later LiqCNA sample (**Fig.5D**). LiqCNA measurements of the growth of the resistant population therefore correlated strongly with disease burden in HGSC patients and so this measure could potentially be useful for tracking ER populations and to guide dose modulations in future trials of adaptive therapy.

**Figure 5.**
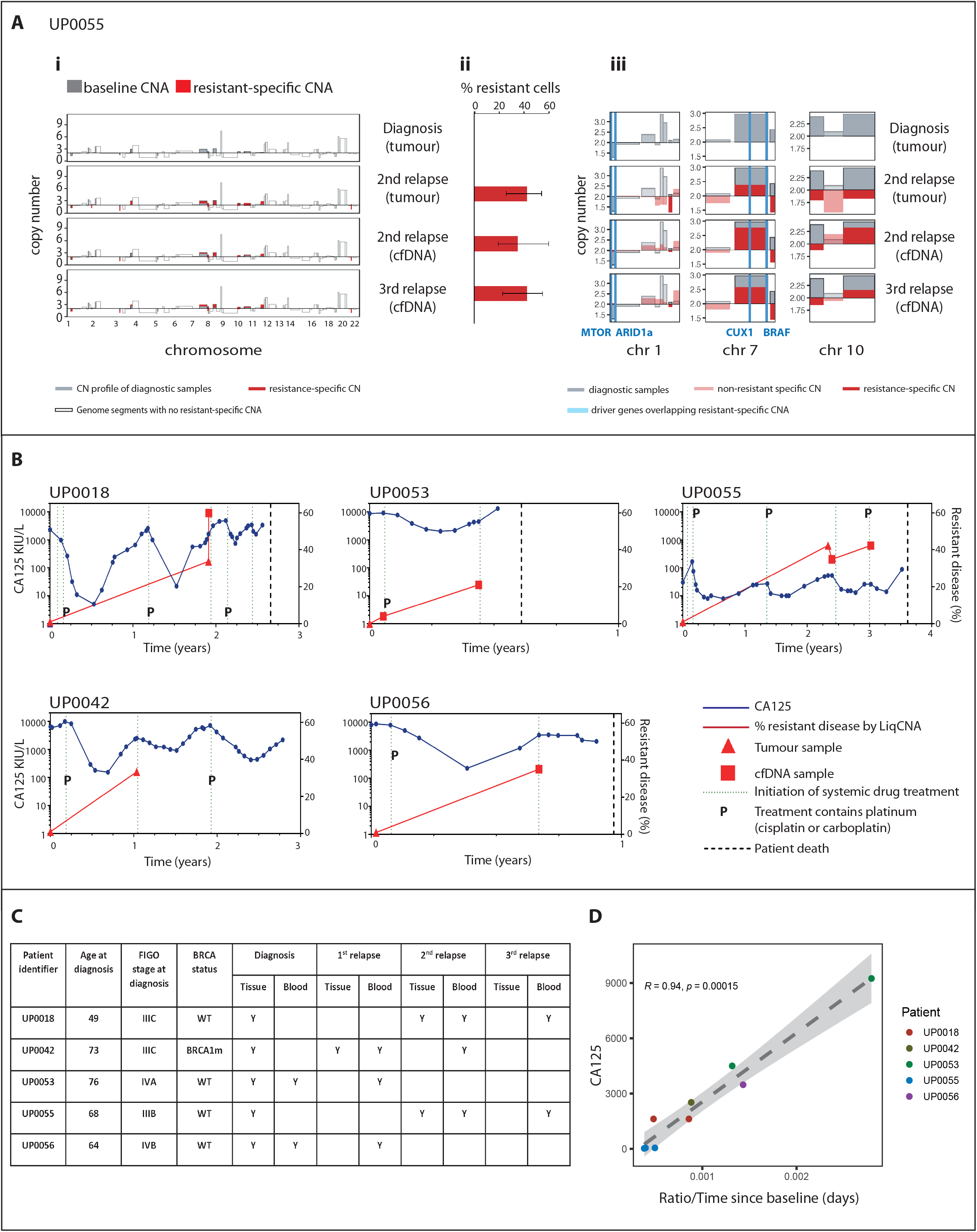
Ai: Tumour purity-corrected copy number profile for each sample obtained from patient UP0055. Grey bars show the copy number (CN) profile of diagnostic samples, and red bars show CN profile of later samples as indicated. Transparent bars show genome segments with no resistant-specific CNAs (i.e. regions with clonally shared CN or with other CNAs) **ii:** Resistant proportion of each sample estimated by LiqCNA. Error bars indicate 95% confidence interval of each estimate. **iii:** Zoomed-in profiles of selected chromosomes with the most prominent/impactful resistant-specific copy number alterations. Driver genes or genes associated with ovarian cancer that overlap with resistant-specific CNA are indicated by blue vertical lines and listed below each graph. **B:** LiqCNA estimation of resistant disease (%, red, right y-axis) and CA125 (kIU/L, blue, left y-axis) plotted over time for five unique patients. Subclonal ratio of tumour and blood samples are shown by triangles and squares respectively. Dotted line indicates start of each new line of chemotherapy, P indicates platinum-containing chemotherapy, dashed line indicates date of patient death. **C:** Tissue and blood samples were collected from 5 patients. Patient age, stage of disease at diagnosis, BRCA mutation status and timings of tissue and blood samples obtained are shown. (BRCA1m = mutation in BRCA1 gene, WT = wild type). **D:** Rate of change in subclonal ratio estimated by LiqCNA in each sequential time point (compared to the diagnostic sample for that patient) plotted against CA125 at the later time point.

## Discussion

Drug therapy in oncology relies on the assumption that greater drug dose results in greater patient benefit and this principle is widely applied throughout cancer drug development. This co-exists with the knowledge that drug resistance is inevitable and that systemic treatment of metastatic cancer is nearly always palliative. AT addresses these conflicting concepts by dynamically tailoring drug dose to the evolution of resistance within individual tumours with the aim of controlling cancer for longer and turning cancer into a chronic disease^4^. AT is based upon the evolutionary principle that resistant phenotypes experience fitness costs in resource-limited environments, due to the cellular resources they require to evade drug therapy^4^. Although this may only represent a small proportion of the energy expenditure of a cell, in the resource-poor conditions typical within the tumour microenvironment, these small changes are sufficient to impair standard cellular processes^31^.

In cancer, the net growth rate (birth-death rates) is a good measure of fitness, and can be estimated by changes in the relative size of sensitive and resistant populations^32^. This appears plausible in HGSC, where sensitivity to first-line platinum-based chemotherapy can be explained by a small, i.e. less fit, resistant population at diagnosis. Furthermore, if resistant cells do have a fitness cost, they would not have expanded in the absence of drug. Equally, the subsequent relapsing/remitting nature implies relative growth and decline of sensitive and resistant populations over time through therapy. Here, we have modelled this situation pre-clinically in a range of HGSC cell lines and *in vivo* models to conclusively show that fitness costs of resistance do exist at low serum conditions. Others have made similar observations following hypoxia in colorectal cancer^33^ and low glucose culture in colorectal^33^ and breast cancer^11^. Although these previous studies demonstrated competition between sensitive and resistant populations, they did not examine the mechanistic basis of the selective decline in resistant populations. We found that this is mediated by apoptosis, presumably when available resources are insufficient to maintain cell viability. This might represent a specific vulnerability of resistant cancer cells that could potentially be exploited in future drug combination strategies.

AT models often focus on competition between sensitive and resistant cancer cells without accounting for other factors that are expected to influence clonal growth rates including spatial constraints and arrangement of cells^34^, highly local levels of shared resource and the contribution of non-cancer cell components of the tumour including vasculature, immune cells and supporting extracellular matrix^35^. Moreover, the role of acquired resistance-conferring mutations, as well as plasticity and phenotype- switching between sensitive and resistant states have rarely been modelled^36^. Our *in vivo* experiments account for some of these factors, although we acknowledge the limitations of the immunodeficient mice used here. Nonetheless, we provide compelling evidence in support of AT, particularly in mice with 100% and 80% sensitive tumours in whom tumour control was significantly extended compared to standard carboplatin treatment with only one of these 10 mice experiencing disease progression during six months of AT. Interestingly, in mice with 100% resistant tumours, standard carboplatin dosing appeared to accelerate tumour growth compared to vehicle and to AT. This observation can be explained by the evolutionary phenomenon of ‘competitive release’, whereby high drug dose eliminates any sensitive cells and facilitates resistant cell growth^37^.

Our AT regimen initially prescribed the same dose of carboplatin as mice receiving standard therapy with the aim of achieving early tumour control. AT dose was then rapidly reduced as tumours shrunk. This regimen meant that by the end of the experiment, mice were receiving repeated, very small carboplatin doses, resulting in higher cumulative drug dose compared to standard treatment. A similar phenomenon has been seen in other pre-clinical AT studies^4, 13^. Our experiment could not determine whether the extended survival with AT was indeed due to evolutionary factors or simply the higher cumulative drug dose received by AT treated mice, although we were encouraged to note that this survival benefit was achieved without excess toxicity. We also note that others have shown that repeated administration of standard therapy failed to achieve the same clinical benefit as AT, implying that the dynamics of AT dosing rather than total drug dose are responsible for the clinical benefit observed^13^. ACTOv will address this by comparing progression-free survival (primary endpoint) as well as toxicity and cumulative carboplatin dose following standard treatment (maximum of six intravenous administrations every three weeks as per standard clinical practice) and AT (maximum 12 administrations, three-weekly) as a secondary trial endpoints.

To date, AT has primarily responded to surrogate measures of total tumour burden, for example changes in the serum tumour marker PSA to direct abiraterone therapy in prostate cancer, and AT in ACTOv will be based on changes in CA125. Other trials have tested ‘intermittent therapy’ in which fixed alternating periods on and off drug are prescribed regardless of tumour response. Examples include the BRAF inhibitor vemurafinib in *BRAF* V600-mutant melanoma^38, 39^; androgen deprivation therapy in metastatic, castration sensitive prostate cancer^40, 41^ and the TKIs sunitinib and pazopanib in clear cell renal cell cancer^42^. Although these trials have generally reported lower toxicity than standard dosing, cancer control has been disappointing and can be explained by the fact that intermittent therapy is not linked to drug response in individual patients over time^43^.

AT is expected to be most effective when dosing regimens relate to, and ideally anticipate, evolving tumour dynamics, however this is challenging to measure, particularly within a clinically relevant time frame. The phase 2 CHRONOS trial in metastatic colorectal cancer (mCRC) provided proof-of-principle that circulating tumour DNA (ctDNA) could be used to guide therapeutic re-challenge with the anti-epidermal growth factor receptor (EGFR) monoclonal antibody, panitumumab, when resistance-associated mutations in *RAS*, *BRAF* and *EGFR* appeared in ctDNA^44^. Unfortunately, resistance to carboplatin is not associated with recurrent point mutations or changes in copy number^25, 27, 28^ and we confirm this in the new patient data presented here. However, we have observed passenger copy number changes that are specific to individual cell lines and HGSC patients^26^. We previously created LiqCNA to measure these changes in multiple sequential samples obtained over time^29^. Here we present novel data to show for the first time that LiqCNA can measure the emergence of resistant ovarian cancer in human patients and that this correlates strongly with changes in CA125. This both endorses our use of CA125 to guide AT in ACTOv and implies that LiqCNA could be a potential biomarker to direct AT in second generation clinical trials. We did however find one example where LiqCNA did not correlate in tissue and blood samples. This could be explained by sampling bias in the tissue sample or potential bias in the blood sample since ctDNA requires shedding from the tumour and so may not be entirely representative of whole tumour population.

In summary, we have shown that AT is significantly more effective than standard carboplatin dosing, achieving long-term control of high grade ovarian cancer. Our finding that sensitive and resistant populations fluctuate through therapy reveals for the first time that the success we observed with AT is indeed due to differences in relative population fitness under and without drug pressure. As a result of these pre-clinical data, carboplatin AT is currently being tested in women with platinum-sensitive, relapsed, high grade ovarian cancer via the ACTOv multicentre, randomised phase 2 clinical trial^24^, which could have important implications for drug dosing in cancer care.

## Methods

### Cell culture and chemotherapy-resistant cell lines

Human HGSOC cell lines OVCAR4 and Cov318 were cultured in DMEM containing 10% fetal bovine serum (FBS) and 1% penicillin/streptomycin. Platinum-resistant cell lines were generated and transfected with dual constructs of Firefly luciferase and either green fluorescent protein (GFP, sensitive ancestral cells) or red fluorescent protein (RFP, evolved resistant cells) as we previously described (CMV-Luciferase (firefly)-2A-GFP/RFP (Puro)^26^.

### Co-culture experiments

Cells were plated in 10% FBS-containing medium at a range of starting ratios of sensitive:resistant cells. Cells were then either maintained in 10% FBS-containing media (high resource) or media was exchanged every 24 hours with fresh 0.5% FBS-containing medium (low resource). In experiments to measure the effect of platinum chemotherapy on cells in co-culture, 50:50 ratios of OVCAR4- GFP:Ov4Cis, cells were plated in 10% FBS and media changed to 0.5% FBS after 24 hours as before. Cisplatin was added to the 0.5% FBS-containing media at a range of doses (1μM, 0.8μM, 0.5 μM and 0.1μM) and administered to the co-cultures on day 6. Exchange with 0.5% FBS-containing media continued every 24 hours for the duration of the experiment. In all cases, co-culture plates were harvested over time and total cell number/μl was recorded (Countess IIR automated cell counter (Life Technologies). Ratio of chemosensitive (GFP) *vs*. chemoresistant (non-GFP) was measured by flow-cytometry (BD LSRFortessa^TM^) and analysed using FlowJo® v8 software. Mathematica v11 and PopDynamics software were used to calculate growth rate and carrying capacity for individual cell populations over time. To test the relationship between the initial seeding ratio and the growth rate of sensitive/resistant cells, we used the lowest sensitive seeding ratio population (5:95 and 15:85 for high and low resource, respectively) to fit the (log) ratio of sensitive to resistant cells over time using a linear model (black lines in **Fig.1E**). The slope of this fitted line measures g=g_s_ – g_r_, the difference in growth rate between sensitive and resistant cells, and is reported in **Fig.1E**. We then aligned the datasets obtained at other seeding ratios so that the first time-point of each dataset fell on the line. We found that all time-points in all datasets closely followed the linear fit, despite these data points not used for fitting, confirming that the dynamics of sensitive to resistant population ratio is independent of the initial seeding ratio.

### Mechanisms of resistance

Cell cycle profiles were acquired by flow cytometry following treatment with propidium iodide (PI) and RNaseA. BV605 Annexin V antibody (BD Horizon^TM^) was used for Annexin V apoptosis assays with 100µM etoposide-treated cells as positive controls. p16 and p21 mRNA was determined by quantitative reverse-transcription PCR (QuantStudio^TM^ 5 Real-Time, Applied Biosystems) normalized to β-actin with 100ng/ml doxorubicin-treated cells as positive controls. (p16: fwd 5′- CAACGCACCGAATAGTTACG-3′; rev 5′-CAGCTCCTCAGCCAGGTC-3′, p21: fwd 5’- GGCAAGAGTGCCTTGA CGAT-3’; rev 5’- CCTCTTGACCTGCTGTGTCG-3’, β-actin: fwd 5’-AGAGCTACGAGCTGCCTGAC-3’; rev 5’- CGTGGATGCCACAGGACT-3’).

### Conditioned media experiment

On day 0, 2.5x10^5^ OVCAR4-GFP or Ov4Cis cells were seeded per well in 6-well plates in 10% FBS-containing medium. After 24 hours the medium was exchanged for 0.5% FBS-containing medium that had been pre-conditioned for 24 hours by either the same cell line, the opposite cell line or a 50:50 OVCAR4-GFP:OV4Cis-RFP co-culture. This same medium was replenished daily. Cells were harvested for up to 11 days and the cell number recorded using a Countess IIR automated cell counter (Life Technologies).

### Animal studies

Experiments were conducted under UK government Home Office project license P1EE3ECB4 following Institutional Review Board approval. Female CD1 nu/nu mice (Charles River Laboratories) were injected subcutaneously with 5x10^6^ cells in 200µl sterile PBS into each flank. Tumours were measured using callipers and volumes calculated using the formula π(short diameter)^2^x(long diameter)/6. Intraperitoneal (IP) tumours were created by IP injection of 5x10^6^ cells in 200µl sterile PBS. Animals were killed via a Schedule 1 method either when the volume of one of their flank tumours had reached 1.44cm^2^, at the end of the experiment or if the project license’s maximum severity was reached. Excised tumours were frozen in liquid nitrogen or immediately fixed in 4% paraformaldehyde.

### GFP and RFP quantification

DNA was extracted using the DNeasy® Blood and Tissue Kit (QIAGEN) following homogenisation and lysis of tissue samples. GFP and RFP DNA was quantified by PCR using QuantStudio^TM^ 5 Real-Time (Applied Biosystems) and normalized to human GAPDH (hGAPDH). (GFP: fwd 5’- GGACGACGGCAACTACAAGA-3’, rev 5’-TTGTACTCCAGCTTGTGCCC-3’; RFP: fwd 5’-TGGTGTAG TCCTCGTTGTGG-3’, rev 5’-ATGAGGCTGAAGCTGAAGGA-3’; hGAPDH: fwd 5’-CCTCACAGTTGCC ATGTAGACC-3’, rev 5’-TCAGTCTGAGGAGAACATACCA-3’).

### Histology

Immunohistochemistry was performed according to standard protocols. Briefly, 4μm serial sections were dewaxed, rehydrated and immersed in 3% hydrogen peroxide for 20 minutes to quench endogenous peroxidase activity. Antigen retrieval was carried out at 95°C for 25 minutes in 10 mM Tris, 1 mM EDTA solution, pH 9.0. After cooling, sections were incubated with blocking buffer (PBS supplemented with 5% goat serum and 1% bovine serum albumin) for 1 hour at room temperature (RT). Primary antibodies were diluted in blocking buffer and applied for 1 hour at RT: rabbit anti-p53 (Cell Signaling Technology, 2527) at 1:200 dilution, rabbit anti-GFP (Cell Signaling Technology, 2956) at 1:75 dilution, anti-Cleaved Caspase-3 (Cell Signaling Technology, 9664) at 1:50 dilution and rabbit anti-luciferase (Abcam ab185924) at 1:200 dilution. Sections were then incubated with a biotinylated anti-rabbit secondary antibody at RT for 45 min, followed by incubation with streptavidin-biotin peroxidase solution at RT for 45 min. Visualization of the first antibody binding was carried out using DAB, according to the manufacturer’s instructions (Vector Labs). Finally, sections were lightly counterstained using Mayer’s haematoxylin and allowed to dry before mounting and digitizing using the Hamamatsu Nanozoomer-XR. Pixels positive for p53 and GFP were quantified using Adobe Photoshop.

### *In vivo* adaptive therapy experiment

Each mouse was randomly allocated to one of three treatment groups when one subcutaneous tumour (target tumour) reached 300mm^3^: 1) Vehicle (200µl of PBS IP every 4 days for 3 doses); 2) Standard therapy (60mg/kg carboplatin IP every 4 days for 3 doses); 3) Adaptive Therapy (AT): one initial dose of 60mg/kg carboplatin IP and then weekly IP carboplatin as follows: target tumour volume changed by ≤20%, administer same dose; target tumour volume increases by >20%, increase dose by 50% (maximum dose 60mg/kg); target tumour volume decreases by >20%, decrease dose by 50%. If the target tumour became smaller than 60mm^3^, treatment was held until target tumour volume was greater than 60mm^3^ again.

### Patient samples

Tumour tissue and blood were obtained from patients with Stage III/IV HGSOC under the ethics of the Barts Gynae Tissue Bank (BGTB) (REC: 20/EE/0193). Blood was collected in 10ml Cell-free DNA BCT® (Streck) tubes and centrifuged within 4 hours of collection (1,200g, 10 minutes, 4^0^C). Plasma supernatant was centrifuged again (4,500g, 10 minutes, 4^0^C) and the supernatant stored at -80^0^C for cell-free DNA extraction and the cell (leucocyte) pellet was stored at -80^0^C for germline DNA extraction.

### DNA extraction and analysis

Leucocyte pellets were treated with red blood cell (RBC) lysis buffer (distilled water containing 155mM NH_4_Cl, 10mM KHCO_3_ and 1mM EDTA) and genomic DNA (gDNA) was extracted (DNeasy® Blood and Tissue Kit (QIAGEN)). DNA was extracted from formalin-fixed paraffin-embedded (FFPE) tissue (High Pure FFPET DNA Isolation Kit (Roche)) following laser capture microdissection in samples with small islands of malignant cells. Genomic and tissue DNA was sonicated into fragments of 150 base pairs using a M220^TM^ Focused-ultrasonicator (Covaris) and library preparation performed using NEBNext® FFPE DNA Repair Mix (New England Biolabs® Inc) and NEBNext® Ultra^TM^ II DNA Library Prep Kit for Illumina® (New England Biolabs® Inc). Cell free DNA (cfDNA) extraction and library preparation was performed on plasma samples (QIAseq^TM^ cfDNA All-in-One Kit (QIAGEN)). Libraries were sequenced with NovaSeq 6000 (Illumina^®^) to an average depth of 0.3x for leucocytes, 0.5x for tissue DNA and 1.9x for cfDNA. Sequencing reads were aligned to genome version hg19 following standard bioinformatic protocols, and copy number profiles were obtained using QDNAseq^45^. Copy number profiles were subsequently analysed using LiqCNA as we previously described^29^ to derive estimates of tumour purity and the resistant population. LiqCNA was run 150 times per patient on a random 75% subsample of genomic segments to derive 95% confidence intervals for each subclonal ratio estimate.

### Statistical analysis

Statistical analysis was performed using GraphPad Prism v7.04. A probability of < 0.05 was considered significant. *=*p* 0.05; **=*p*<0.001; ***=*p*<0.0001. A two-tailed paired *t-*test was used unless otherwise stated. The Pearson correlation coefficient (R) was calculated to measure linear correlation between two sets of data with R^2^ indicating the goodness of fit of a model.

## Acknowledgements

We thank staff in the core facilities at Barts Cancer Institute including FACS, the Barts Gynae Tissue Bank and the animal technician service. We thank Dr Jacqueline McDermott for help with slide interpretation and thank the patients and clinical teams who contributed to this work.

