## Supplemental Figures and Legends for "Adaptive therapy achieves long-term control of chemotherapy resistance in high grade ovarian cancer"

### Supplementary Figure Legends

**S1:**

Abundance of sensitive (OVCAR4-GFP) and resistant (Ov4Cis) cells co-cultured over time in high resource conditions (10% FBS) at a range of starting ratios. Green=OVCAR4-GFP, red=Ov4Cis, mean±st.d, *n*=3 technical replicates).

**S2:**

Sensitive and resistant cell abundance over time for co-culture different seeding ratios in high resource conditions (10% FBS). $g_S$ (exponential growth rate of sensitive cells) and $g_R$ (exponential growth rate of resistant cells) are shown for each seeding ratio.

**S3:**

Abundance of three different sensitive (green) and resistant cell pairs (red) over time when grown as co-cultures in low resource conditions (0.5% FBS exchanged daily) at three starting ratios (85% sensitive, 50% sensitive and 15% sensitive). Solid lines indicate observed cell growth, dashed lines indicate predicted cell growth based on 100% monoculture experiments.

**S4:**

**A:** Doxorubicin-induced expression of p16 and p21 in OVCAR4-GFP and Ov4Cis cells by reverse transcription qPCR (mean±st.d, *n*=3, **p<0.01, unpaired *t*-test). **B:** OVCAR4-GFP and Ov4Cis cells were grown as mono-culture and as 85:15 S:R and 50:50 S:R co-cultures. Co-culture samples were sorted into GFP-positive and GFP-negative populations by flow cytometry. Expression of p16 and p21 by reverse transcription qPCR in each population is shown (mean±st.d, *n*=3). Horizontal dotted lines indicate a 2-fold and 0.5-fold increase in gene expression.

**S5:**

OVCAR4-GFP (sensitive, green) and Ov4Cis-RFP (resistant, red) were co-cultured at different starting ratios (85:15, 50:50 and 15:85) in low resource conditions (media exchanged with fresh 0.5% FBS-containing media every 24 hours). Cells were harvested over time and GFP-/RFP-positive cells quantified by flow cytometry (mean±st.d, *n*=3 technical replicates).

**S6:**

**A:** Target tumour volume over time in the two mice with 80:20 sensitive:resistant tumours, who received Adaptive Therapy and were culled before experimental endpoint. Dotted line indicates initiation of Adaptive Therapy at Time=0, X indicates date at which each mouse was culled. **B:** Immunohistochemistry of resected tumours from Mouse 1 (left) and Mouse 2 (right). p53 indicates tumour cells and Firefly luciferase indicates the resistant population.

**S7:**

**A-Di**: Tumour purity-corrected copy number profile for each sample obtained from patients UP0018 (**A**), UP0042 (**B**), UP0053 (**C**) and UP0056 (**D**). Grey bars show the copy number of each segment in the baseline diagnostic tumour biopsy sample and red bars show CN profile of later samples as indicated. **A-Dii:** Resistant proportion of each sample estimated by LiqCNA. Error bars indicate 95% confidence of each estimate. **A-Diii:** Zoomed-in profiles of selected chromosomes with the most prominent/impactful resistant-specific copy number alterations. Driver genes or genes associated with ovarian cancer that overlap with resistant-specific CNA are indicated by blue vertical lines and listed below each graph.

S1

**OVCAR4-GFP : Ov4Cis**

(High Resource = 10% FBS)

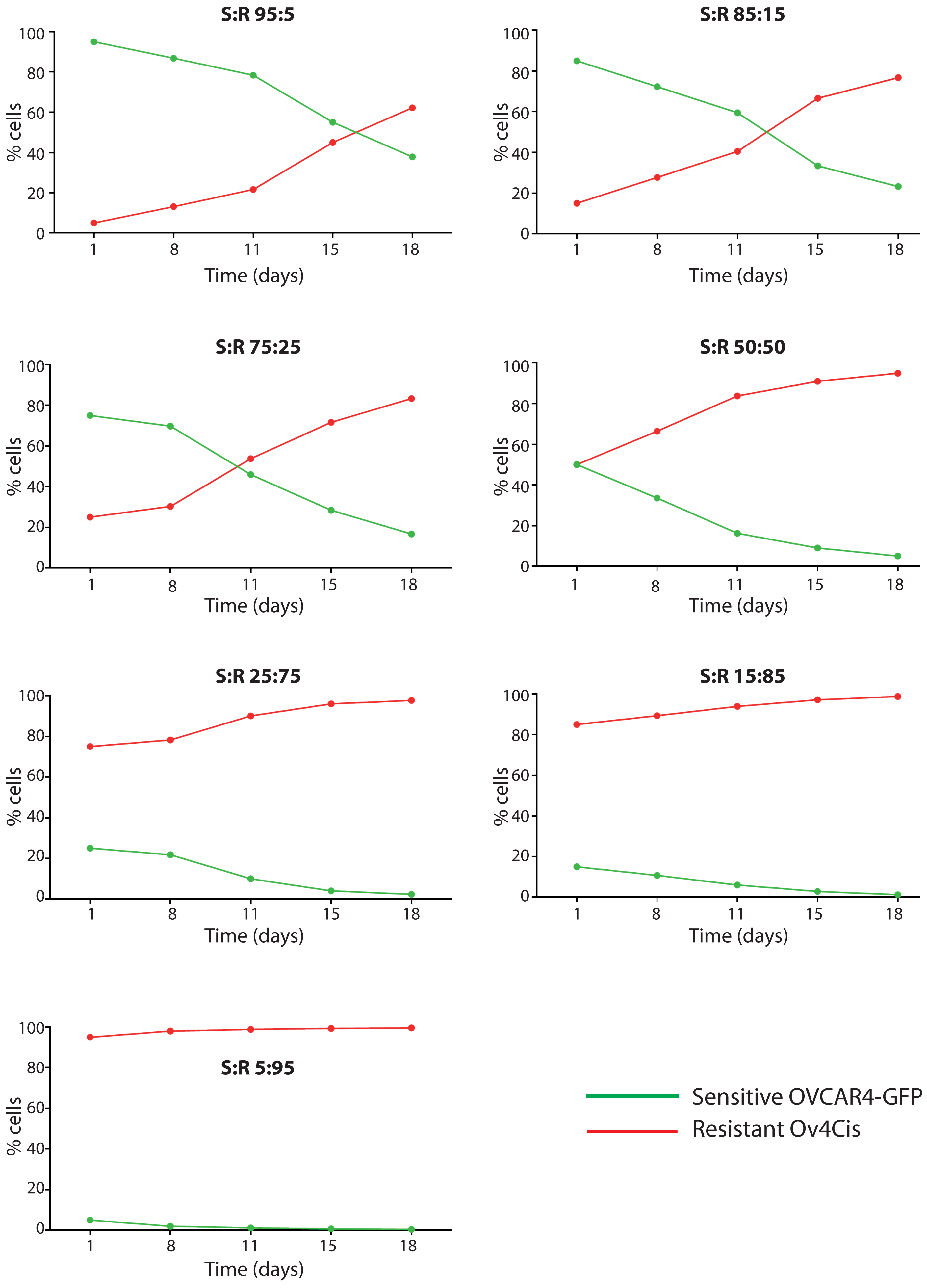

S2

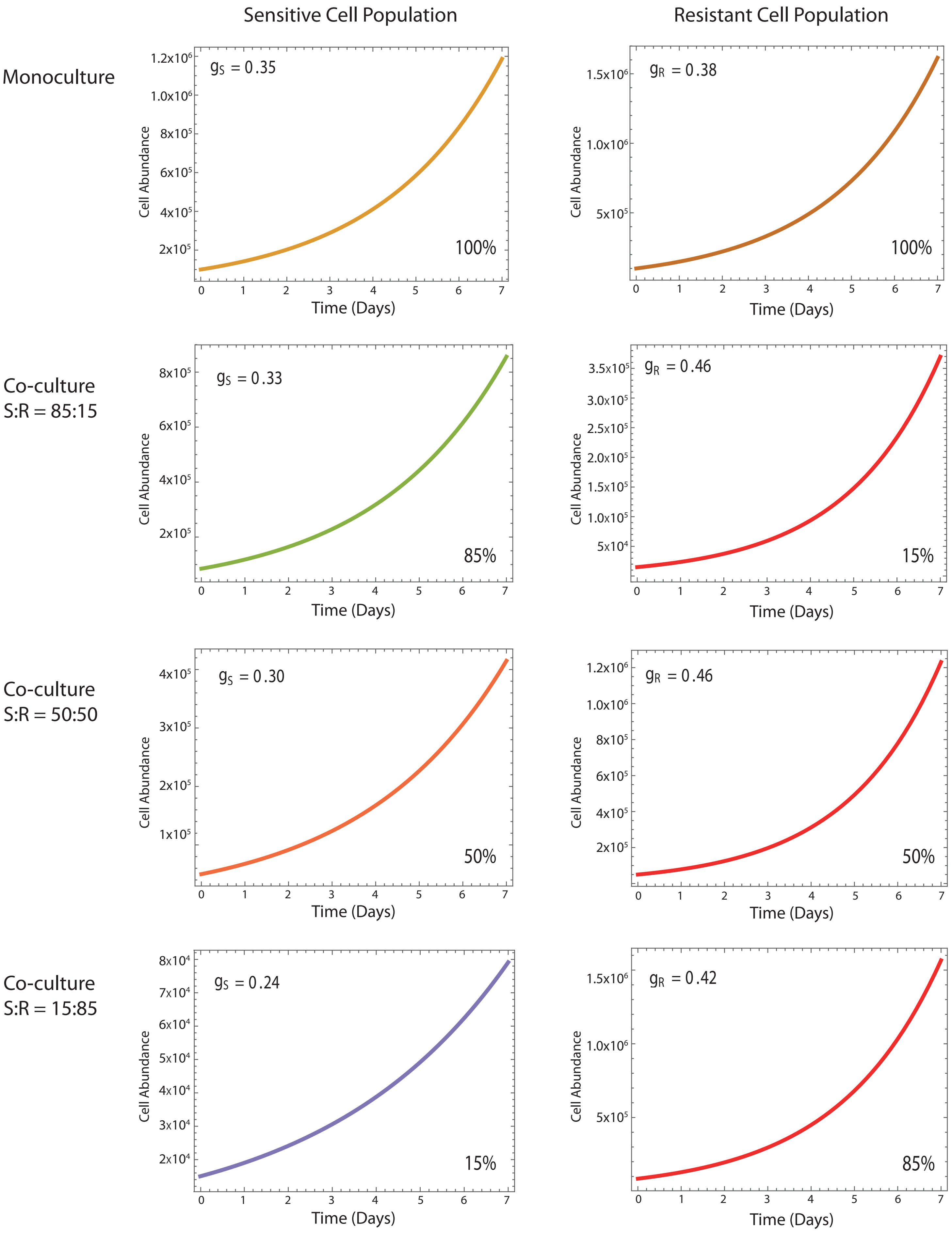

Sensitive Cell Population
Resistant Cell Population
Monoculture
Co-culture S:R = 85:15
Co-culture S:R = 50:50
Co-culture S:R = 15:85
$g_S = 0.35$
$g_R = 0.38$
$g_S = 0.33$
$g_R = 0.46$
$g_S = 0.30$
$g_R = 0.46$
$g_S = 0.24$
$g_R = 0.42$
100%
100%
85%
15%
50%
50%
15%
85%
Cell Abundance
Time (Days)

S3

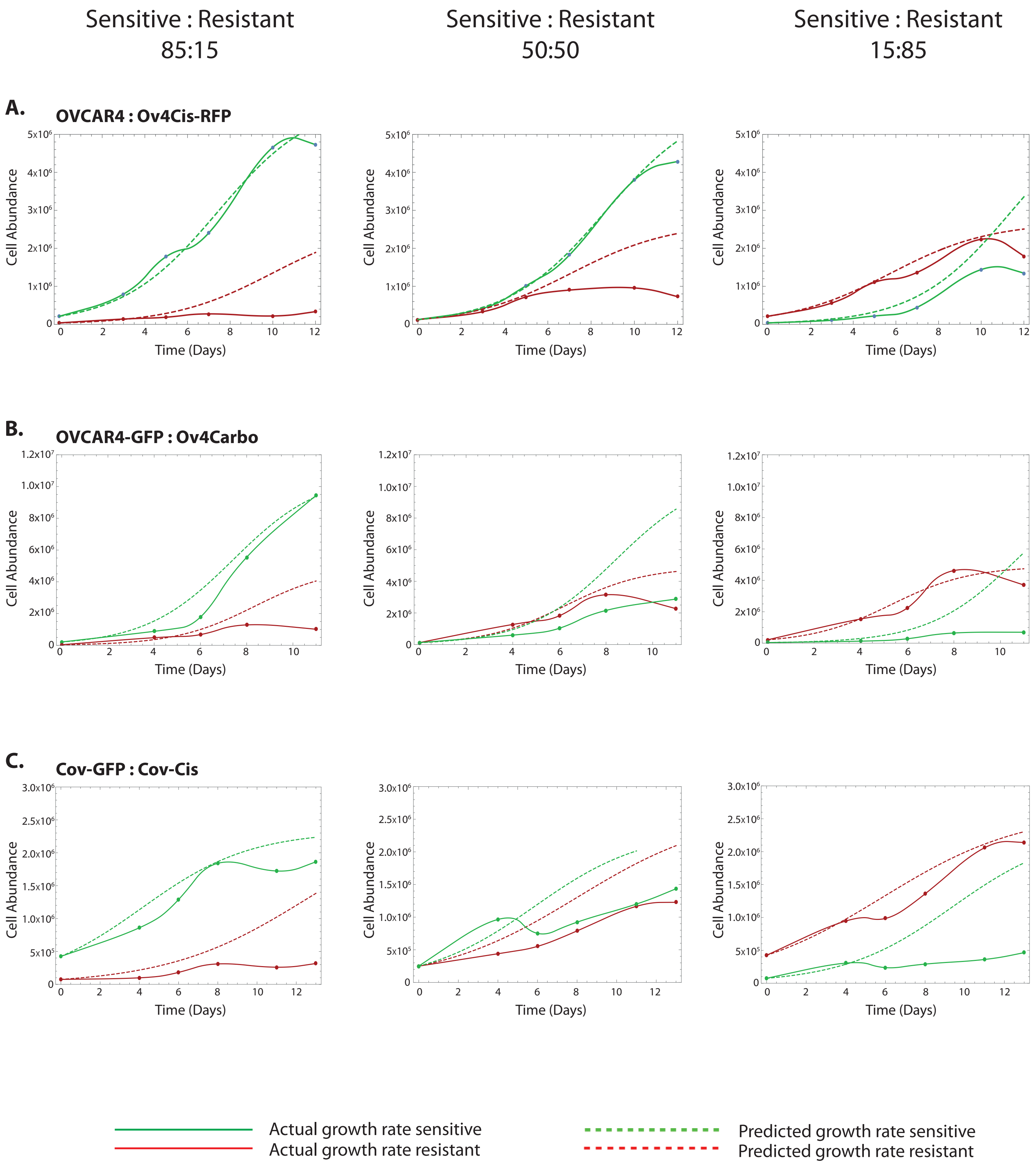
Sensitive : Resistant
85:15
Sensitive : Resistant
50:50
Sensitive : Resistant
15:85
A.
OVCAR4 : Ov4Cis-RFP
B.
OVCAR4-GFP : Ov4Carbo
C.
Cov-GFP : Cov-Cis
Cell Abundance
Time (Days)
Actual growth rate sensitive
Actual growth rate resistant
Predicted growth rate sensitive
Predicted growth rate resistant

S4

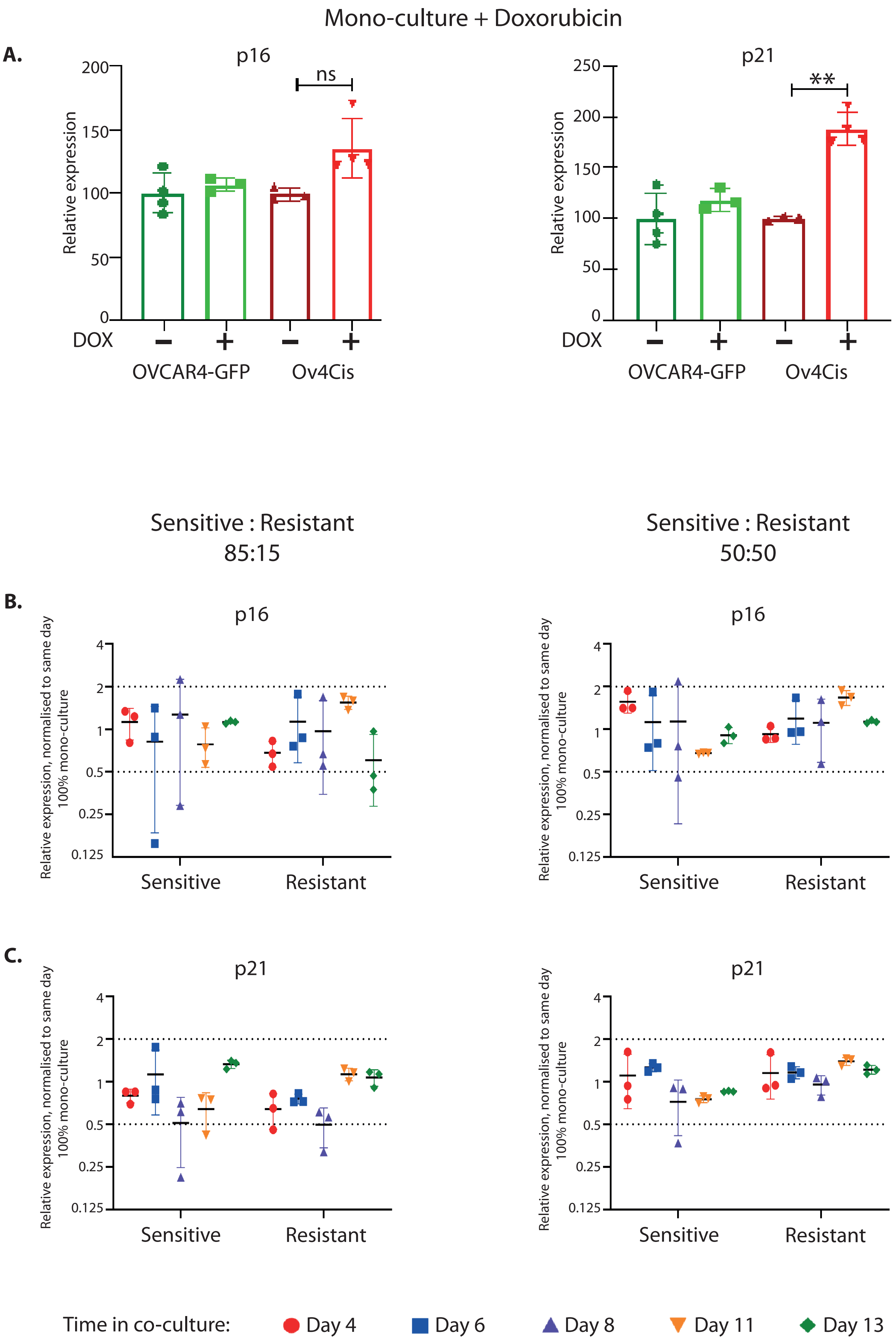

Mono-culture + Doxorubicin
A.
p16
ns
Relative expression
200
150
100
50
0
DOX
OVCAR4-GFP
Ov4Cis
p21
**
250
Sensitive : Resistant
85:15
50:50
B.
p16
Relative expression, normalised to same day
100% mono-culture
4
2
1
0.5
0.25
0.125
Sensitive
Resistant
C.
p21
Time in co-culture:
Day 4
Day 6
Day 8
Day 11
Day 13

S5

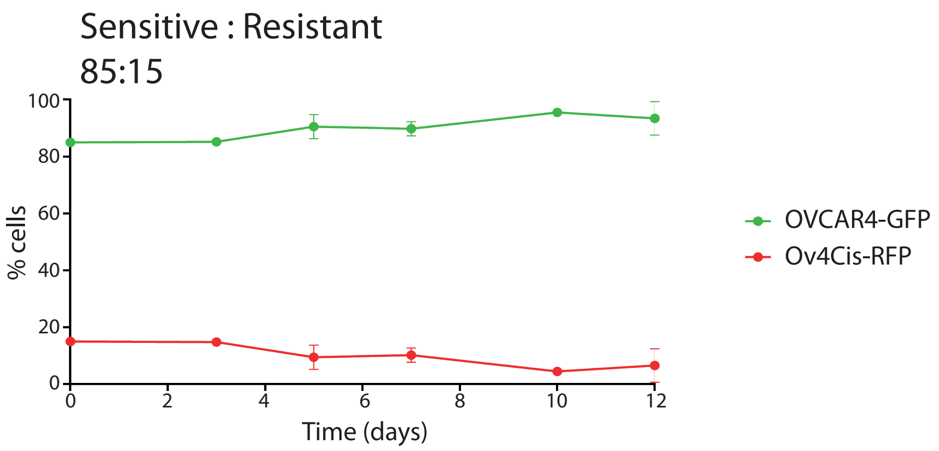
Sensitive : Resistant
85:15
100
80
60
40
20
0
% cells
0
2
4
6
8
10
12
Time (days)
OVCAR4-GFP
Ov4Cis-RFP

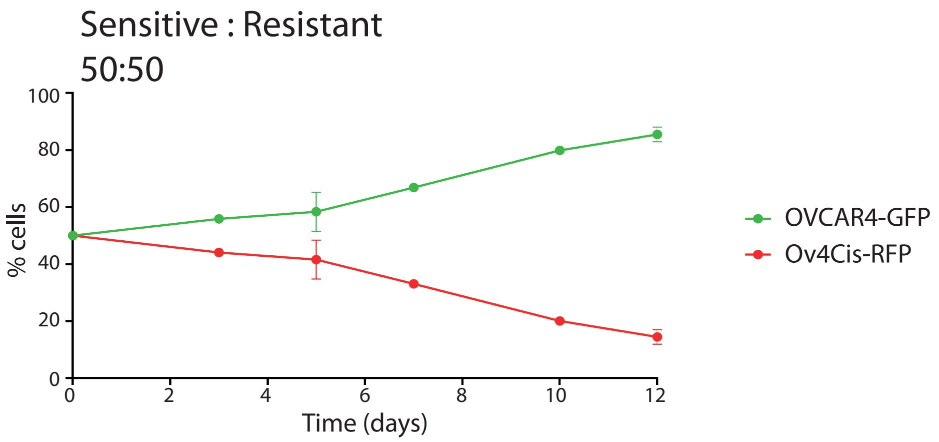
Sensitive : Resistant
50:50
100
80
60
40
20
0
% cells
0
2
4
6
8
10
12
Time (days)
OVCAR4-GFP
Ov4Cis-RFP

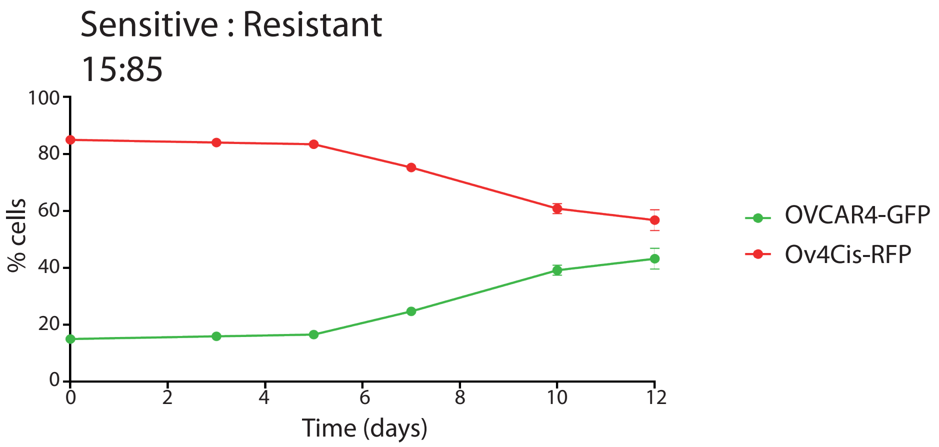
Sensitive : Resistant
15:85
100
80
60
40
20
0
% cells
0
2
4
6
8
10
12
Time (days)
OVCAR4-GFP
Ov4Cis-RFP

S6

**80 : 20**
**Sensitive : Resistant**

**A.**

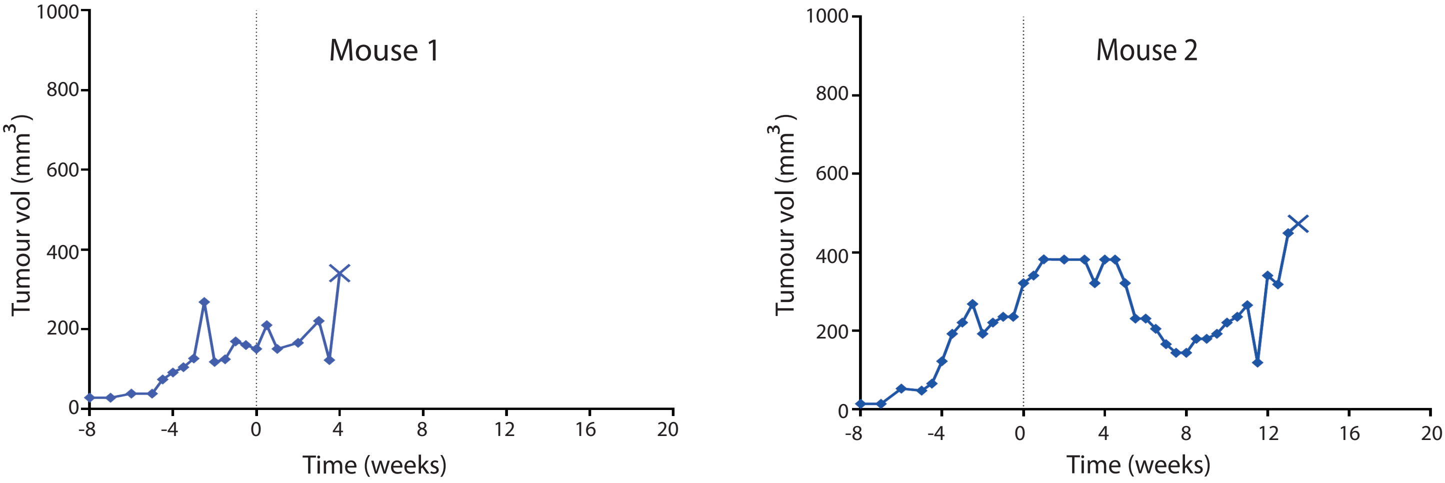


Dotted line = initiation of Adaptive Therapy

X = mouse culled

**B.**

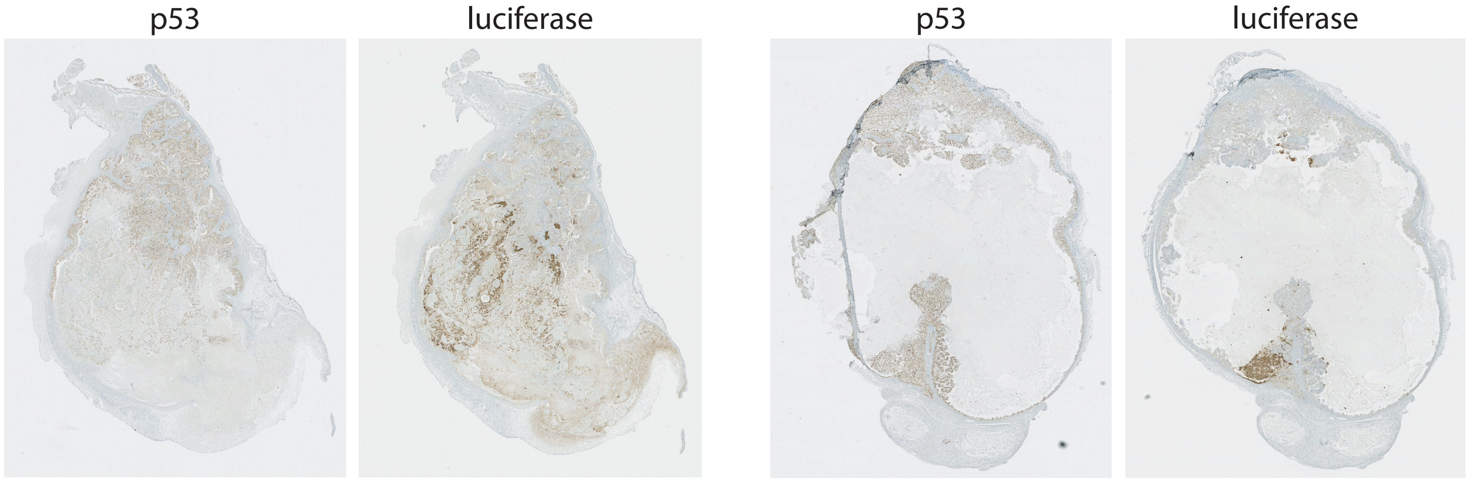

# S7

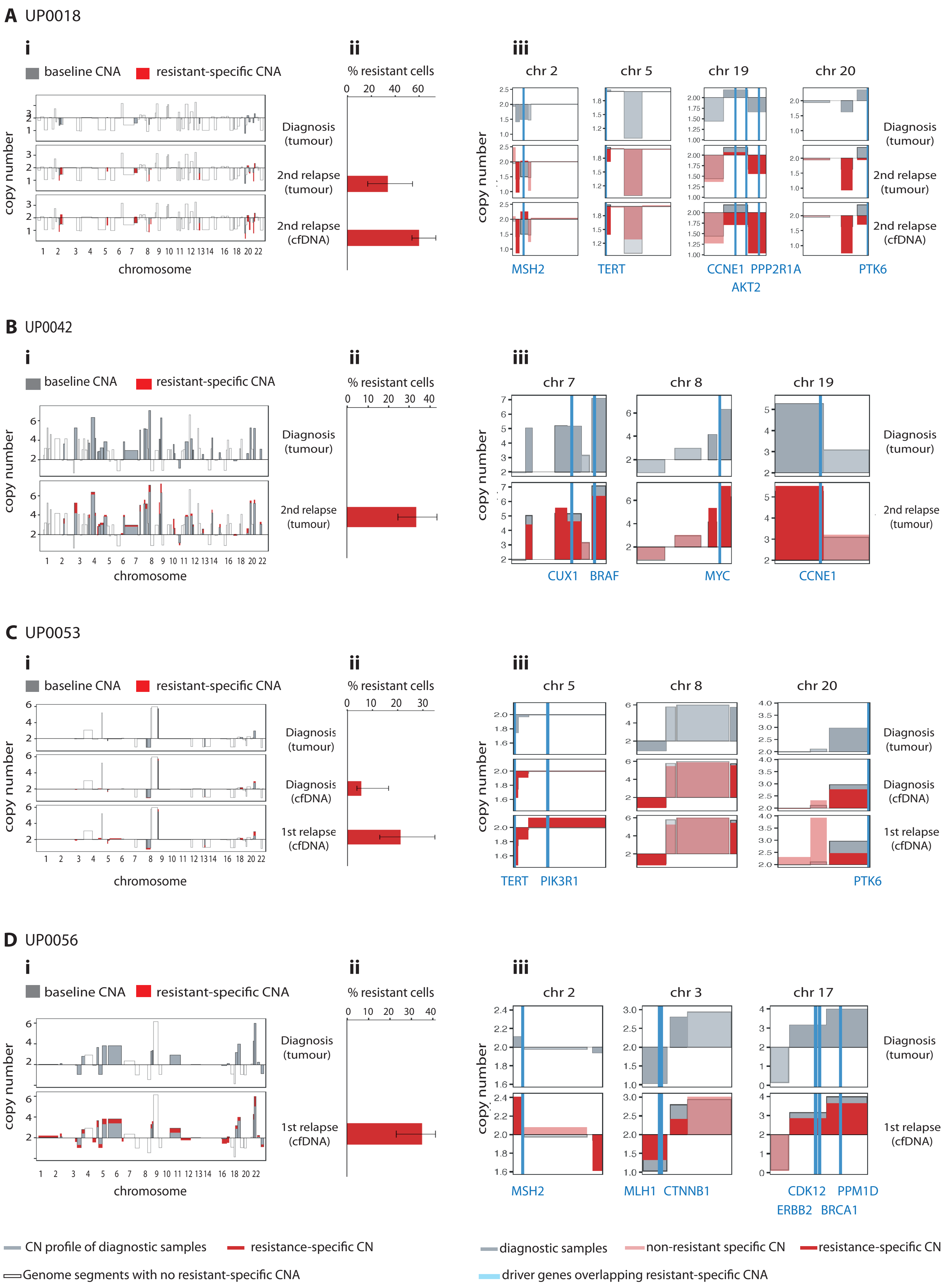
A UP0018
i
baseline CNA
resistant-specific CNA
copy number
chromosome
ii
% resistant cells
Diagnosis (tumour)
2nd relapse (tumour)
2nd relapse (cfDNA)
iii
chr 2
chr 5
chr 19
chr 20
MSH2
TERT
CCNE1
PPP2R1A
AKT2
PTK6
B UP0042
i
baseline CNA
resistant-specific CNA
copy number
chromosome
ii
% resistant cells
Diagnosis (tumour)
2nd relapse (tumour)
iii
chr 7
chr 8
chr 19
CUX1
BRAF
MYC
CCNE1
C UP0053
i
baseline CNA
resistant-specific CNA
copy number
chromosome
ii
% resistant cells
Diagnosis (tumour)
Diagnosis (cfDNA)
1st relapse (cfDNA)
iii
chr 5
chr 8
chr 20
TERT
PIK3R1
PTK6
D UP0056
i
baseline CNA
resistant-specific CNA
copy number
chromosome
ii
% resistant cells
Diagnosis (tumour)
1st relapse (cfDNA)
iii
chr 2
chr 3
chr 17
MSH2
MLH1
CTNNB1
CDK12
PPM1D
ERBB2
BRCA1
CN profile of diagnostic samples
resistance-specific CN
Genome segments with no resistant-specific CNA
diagnostic samples
non-resistant specific CN
resistance-specific CN
driver genes overlapping resistant-specific CNA
